# Training interoceptive awareness with real-time haptic vs. visual heartbeat feedback

**DOI:** 10.1101/2024.01.12.575196

**Authors:** Olga Dobrushina, Yossi Tamim, Iddo Yehoshua Wald, Amber Maimon, Amir Amedi

## Abstract

The perception of signals from within the body, known as interoception, is increasingly recognized as a prerequisite for physical and mental health. The study is dedicated to the development of effective technological approaches for enhancing interoceptive awareness. We provide evidence of the effectiveness and practical feasibility of a novel real-time haptic heartbeat supplementation technology combining principles of biofeedback and sensory enhancement. In a randomized controlled study, we applied the developed naturalistic haptic feedback on a group of 30 adults, while another group of 30 adults received more traditional real-time visual heartbeat feedback. A single session of haptic, but not visual heartbeat feedback resulted in increased interoceptive accuracy and confidence, as measured by the heart rate discrimination task, and in a shift of attention towards the body. Participants rated the developed technology as more helpful and pleasant than the visual feedback, thus indicating high user satisfaction. The study highlights the importance of matching sensory characteristics of the feedback to the natural bodily prototype. Our work suggests that real-time haptic feedback might be a superior approach to strengthen mind-body interaction in interventions for physical and mental health.

**Highlights:**

- We developed a naturalistic real-time haptic feedback system for enhancing cardiac interoception.
- The intervention resulted in higher interoceptive accuracy and confidence, and in a shift of attention towards the body.
- Haptic feedback outperformed traditional visual feedback in efficacy and user satisfaction.
- Results of the study indicate the importance of feedback sensory characteristics in mind-body technologies.

## 1. Introduction

An emergent direction of contemporary neurotech is developing approaches effectively targeting interoception. Interoception is the amalgamation of processes allowing for the perception, interpretation, and integration of signals originating from within the body (Khalsa et al., 2018). Interoception is now accepted to be of great relevance to numerous domains. With respect to emotion, it has been tied to social cognition, affective regulation and emotional processing (Dobrushina, Dobrynina, et al., 2020; Gao et al., 2019; Terasawa et al., 2013; Zaki et al., 2012). In the cognitive domain, interoception has been correlated with self-consciousness, perception of time, memory, decision making, among other cognitive processes (Sadeghi et al., 2023; Tsakiris & Critchley, 2016; N. S. Werner et al., 2009). Interoception is also becoming ever more significant in the realm of mental health, as abnormal or reduced interoceptive processing is associated with various neurological and psychological conditions and exacerbations of their symptoms (Bonaz et al., 2021; Brewer et al., 2021; Nord & Garfinkel, 2022), among them depression and anxiety (Paulus & Stein, 2010), eating disorders (Khalsa et al., 2022), Alzheimer’s disease and other neurodegenerative disorders (Dobrushina, Arina, et al., 2020; Sun et al., 2022). Interoceptive abilities have been correlated with altruistic and prosocial tendencies (Contreras-Huerta et al., 2023; Piech et al., 2017), and lower symptom severity in sufferers of chronic conditions (Locatelli et al., 2023).

In psychology, the most employed method for improving or enhancing interoception is mindfulness training. Though considered to be effective (Bornemann & Singer, 2017), mindfulness practice is notoriously difficult for certain individuals, such as people with low baseline interoceptive abilities, who are in essence those who could benefit from it the most. And even people without such difficulties find it challenging to start and maintain complex mindfulness practices. Another limitation of mindfulness is that while it effectively targets awareness to breathing, low intensity of other interoceptive signals such as heartbeats make it challenging to focus on them in a natural way. Most evidence on the significance of interoceptive abilities for emotion, social efficacy, cognition, and mental health comes from the field of cardioception, warranting the development of technological aids to support heartbeat awareness (Meyerholz et al., 2019).

Numerous previous studies have utilized classical biofeedback for cardioceptive training. Ashton et al. (1979) were among the first to provide visual feedback to participants during a 2-alternative forced choice task designed to train perceptual accuracy with respect to one’s heart rate (not specific heartbeats or real-time interbeat interval variability), showing that this accuracy can be increased following such training (Ashton et al., 1979). This was later replicated by Grigg and Ashton (1982), who extended the procedure to include an exteroceptive form of biofeedback presented to one experiment group by means of a flashing light (Grigg & Ashton, 1982). Schandry & Weitkunat (1990), provided auditory feedback in real time to the pace of the heart, which either did or didn’t taper off, while the participants responded to the heartbeats by button press (Schandry & Weitkunat, 1990). They showed improvement in cardiac perception, and enhancement in the participants’ neural response associated with heartbeats (heartbeat evoked potential). Schaefer et al. (2014) later employed a heartbeat training procedure, in which visual biofeedback of the participants’ beating heart was presented in the form of a red heart animation (Schaefer et al., 2014). Using this procedure, they showed improved heartbeat perception, but moreover, their findings indicated a link between lower cardioceptive accuracy and greater symptom severity in sufferers of these disorders. Several studies have since employed similar cardiac biofeedback methods based on visual feedback by way of a symbolic representation of a heart, such as Meyerholz et al. (2019), Schillings et al. (2022) showing increased cardioceptive awareness following the training (Meyerholz et al., 2019; Schillings et al., 2022). Besides improving interoception, cardioceptive training has been shown to decrease anxiety in autistic adults (Quadt et al., 2021).

Considering the efficacy of classical biofeedback in improving cardioception, we hypothesized that the training could be even more effective if it were to additionally incorporate principles of sensory augmentation. Previous findings coming from our and other groups indicate that the unisensory perceptual capabilities such as vision or hearing can be enhanced through the addition of supplementary, simultaneous feedback through additional senses (Abboud et al., 2014; Auvray et al., 2007; Bach-Y-Rita, 2004; Chebat et al., 2007; Cieśla et al., 2019, 2022; Gori et al., 2014; Haigh et al., 2013; Ptito, 2005). Furthermore, training with these sensory substitution and augmentation devices can induce fundamental organizational and developmental plastic changes in the brain (Abboud et al., 2014; Aggius-Vella et al., 2023; Striem-Amit & Amedi, 2014). Recently, we have shown that employing an immersive sensory experience that responds in real time to one’s physiology significantly increases one’s sense of embodiment (Wald et al., 2023).

Building on this and considering the known links between external and internal body perception (Suzuki et al., 2013; Tsakiris et al., 2011), we supposed that the natural low-intensity signals from the beating heart might be effectively supplemented by congruent haptic stimulation applied to the chest area. Relying on the principles of sensory substitution and augmentation, we put emphasis on accurate depiction of both heart rhythm and pulse wave. Thus, we developed a novel approach employing a real-time haptic biofeedback mimicking the natural sensations of one’s own heartbeats. In this study, we test whether such body related feedback will be similarly (or more or less) effective than the previously utilized non-bodily (e.g. visual) feedback.

## 2. Methods

### 2.1 Participants and study design

The study sample consisted of 60 participants aged 28±9.8, 41 females, 18 males and 1 participant who preferred not to disclose the gender. Participants were recruited through the SONA participant pool management system of Reichman University students, and through the social networks of the researchers according to the following inclusion criteria: participants were required to be 18 years old or above, with normal or corrected-to-normal vision, and fluent English speakers.

The study duration was approximately 1.5 hours. First, participants were given brief explanations. They were told that the study was aimed at testing different ways of improving interoception, that they will receive interoceptive training preceded and followed by interoceptive abilities test, and that the type of the training will depend on the order of randomization. To avoid bias, participants were not made aware of the specific interest in the effect of haptic feedback, and thus, could not discern whether they were in the experimental or control group.

The central portion of the experiment consisted of a heartbeat perception training session. This 12 min session included either haptic or visual feedback of the participants’ heartbeats. The type of the training (haptic or visual) was determined on a 1:1 randomization rate. Before and after the training participants performed a heart rate discrimination task and filled out the state section of the Spielberger state-trait anxiety questionnaire (Spielberger et al., 1983). During the training, participants rated the direction of attention (“sensation vs. thoughts”) three times: after the instruction phase, immediately before the first training run, in the middle of the training and immediately after the last training run.

The study was approved by the Reichman University Institutional Review Board (ethical clearance number P_2023030), and all participants gave written informed consent. The study was performed in accordance with the Declaration of Helsinki and all other relevant guidelines and regulations.

### 2.2 Haptic feedback of the heartbeats

The experiment setup included a laptop with an additional screen for participants, a custom built haptic feedback device, and a Plux biosignals sensor with single use ECG electrodes (see Fig. 1). During the training, participants were seated at a table in a quiet room facing a monitor. Haptic feedback of the heartbeats was provided using a Woojer vibration speaker located in the anterior middle section of the chest, above the heart area, and fixed with an elastic strap. The device was connected to the computer audio port. A Plux biosignals sensor acquired 500 Hz ECG signals from electrodes positioned in the second lead; the signal was received through the Bluetooth port into the OpenSignals software and sent out via lab streaming layer (LSL).

**Figure 1.**
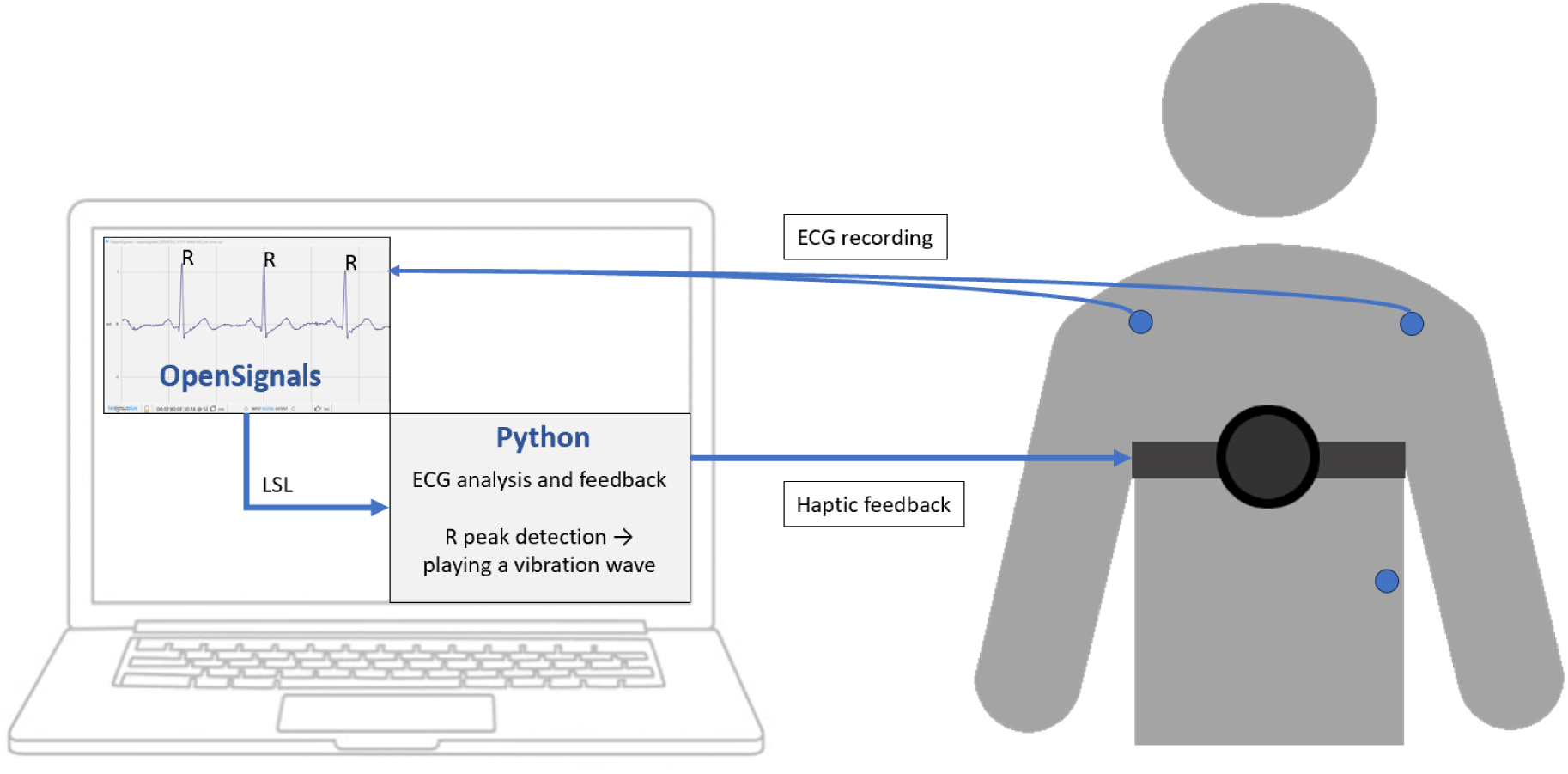
Real-time haptic feedback setup. *Note.* A vibration device located over the heart area is used to provide haptic stimulation mimicking the sensations from the beating heart, while electrocardiographic (ECG) signal is recorded to achieve precise synchronization. The stimulation begins when ECG R-peaks is detected, i.e. at the beginning of systole, and the vibration waveform mimics the dynamics of pressure rise during blood ejection. The setup is operated using a specially developed Python script.

A custom Python script was used to acquire real-time ECG signal through LSL, detect ECG R-peaks (corresponding to the heartbeats), present instructions and provide haptic stimulation for each heartbeat. To identify R-peaks, a 30 sec baseline ECG fragment was recorded for each participant (before the initiation of stimulation), analyzed using the Neurokit2 library (https://github.com/neuropsychology/NeuroKit; Makowski et al., 2021) to detect ECG peaks, and then the R-peak threshold was defined as threshold=noisePeak + 0.25*(signalPeak-noisePeak). SignalPeak is the 25% percentile for R-peaks height and noisePeak is the 75% percentile for T-peaks height in the recorded ECG fragment. This algorithm follows the principles of the classical Pan and Tompkins approach (Tompkins, 1985). During the stimulation, the moment was classified as “peak detection” if the ECG signal exceeded the threshold, and no peak was detected during the previous 0.3 sec. The “latent window” of 0.3 sec was defined on the base of expected heart rate of no more than 200 beats per minute (60 sec/200=0.3 sec). The quality of ECG data was controlled manually after electrode placement and offline on the base of ECG recordings from the training.

Detection of an R-peak initialized playback of an audio stimuli sent to the Woojer device. The audio stimulius was designed with the goal of mimicking the natural sensations from the systolic pressure rise in the heart ventricles. It represented a 38 Hz vibration wave with the amplitude shaped by a triangular ascending-descending function with start, peak and end points corresponding to the left ventricular pressure systolic dynamics: start at R-peak detection, peak 200 ms and end 400 ms after. The baseline frequency of 38 Hz was experimentally identified as the optimal balance between higher frequencies that felt too superficial, at the level of the skin rather than inside the chest, and lower frequencies that created sensations which were determined to be too weak. Audio stimuli was generated in Max MSP. Stimuli were presented using the “Psychopy” toolbox with the ‘PTB’ audio library.

The training included the following steps:

1. Recording 30 sec ECG fragment and R-peak threshold definition.
2. First attention direction rating (see below).
3. Individual determination of the vibration amplitude resulting in comfortable sensations (clearly felt, but gentle, not disturbing) — comfortable amplitude.
4. Individual determination of the minimal vibration amplitude when the sensations can be still felt (sensory threshold) — minimal amplitude.
5. Six runs of training with “fading-out” haptic feedback, 2 min each, with second attention direction rating in the middle of the training.
6. Third attention direction rating.

During each training run, for 95 sec the amplitude of vibration was linearly decreased starting from the comfortable level and ending at the minimal level, for 5 sec it was kept at the minimal level, for 10 sec it was reduced from minimal level to zero, and then left at zero for the remaining 10 sec. The participant received the following instruction: “During the training please focus attention in the center of the chest and attend to your heartbeats. The device will provide a vibration for each heartbeat to help you recognize them. Keep your eyes open.” The remaining time in each run was indicated on the screen by a progress bar.

### 2.3 Control task: visual feedback

The control task consisted of visual feedback of the participants’ heartbeats and followed the same sequence as the haptic feedback task. Vibration amplitude adjustment steps were omitted, and instead of haptic feedback the participants were provided with an image of a heart on the screen, enlarging at 200 ms and returning to the regular size at 400 ms after the R-peak, resulting in an animation of a beating heart synchronized with the real heart’s beats. The last 20 seconds of each run the heart on the screen disappeared. The remaining time in each run was indicated on the screen by a progress bar.

### 2.4 Direction of attention rating

Direction of attention rating was performed using a visual single-item scale. Participants were asked to choose (by clicking) one of the 5 images describing their state on a continuum from “being in sensations” to “being in thoughts” (see Fig. 2). The images were designed by the authors based on the self-assessment manikin “heart vs. mind” (Imbir, 2016). Ratings were assigned from 1 (“I am in my sensations”) to 5 (“I am in my thoughts”).

**Figure 2.**
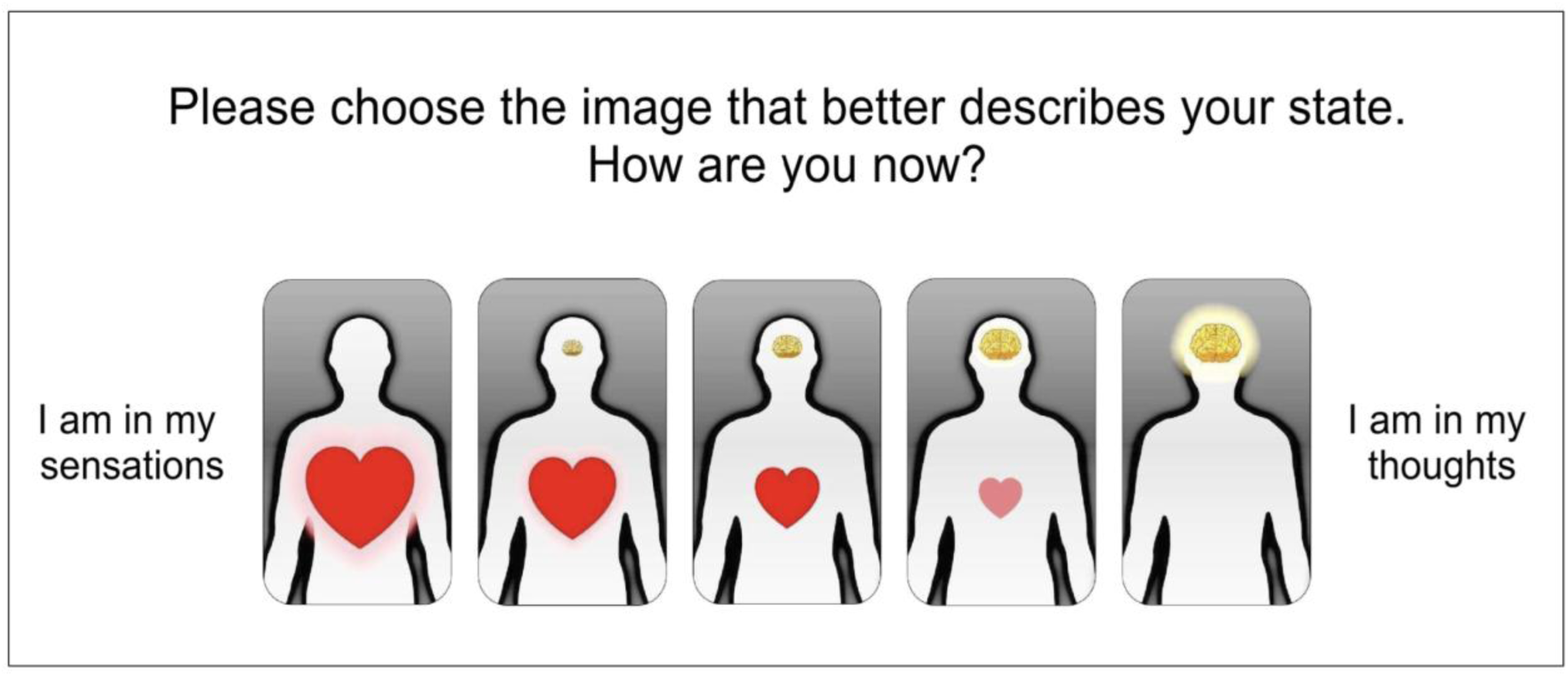
Attention rating scale. *Note:* Specially designed single-item attention rating scale was provided to the participant after the instruction, in the middle of the training and after the training.

### 2.5 Heart rate discrimination task

To test interoceptive abilities, we utilized the heart rate discrimination task previously developed and validated by Legrand et al. (2022). This heart rate discrimination task aims to measure different aspects of cardioception. The task included thirty interoceptive and thirty exteroceptive trials. During interoceptive trials, participants first attend to their cardiac sensations for 5 seconds while their heart rate was monitored via electrocardiography (ECG). Following this “heart listening” phase, participants heard a sequence of five auditory tones and needed to decide whether these tones were faster or slower than their estimated heart rate by pressing the left or right mouse button. After the decision participants rated their confidence on a scale from 0 to 100. The task also included an exteroceptive control condition, where participants compared two sequences of auditory tones instead of comparing an auditory tone to their heart rate.

The task utilized a staircase procedure to adaptively change the frequency of the second set of tones based on the participant’s responses. For the interoceptive modality, the frequency of the feedback tones was adjusted to match the participants’ recorded heart rate, with an additional value denoted as Δ-BPM. The objective was to identify the point of subjective equality (threshold), at which participants are equally likely to judge the feedback sequence as “Faster” or “Slower” than their heart rate. In exteroceptive trials, the frequency of the second set of tones was manipulated relative to a randomly selected reference frequency, following similar principles to the interoceptive condition. In both interoceptive and exteroceptive trials Δ-BPM was adjusted using an adaptive Bayesian approach to estimate the threshold and slope. Interoceptive threshold correlates to the classical interoceptive accuracy measure (Legrand et al., 2022).

The task was obtained from the repository provided by the developers of the test (https://github.com/embodied-computation-group/Cardioception), modified to work with ECG equipment instead of pulse oximeter, and presented using a Python code based on the Psychopy library (https://www.psychopy.org/). ECG was collected using Plux biosignal sensors and single use electrodes positioned in the second lead; the signal was received through Bluetooth port into the OpenSignals software and transferred to Python in real time using LSL. ECG R-peak detection was performed using the Neurokit2 library. The quality of ECG data was controlled manually after electrode placements and offline on the base of ECG recordings from the training.

### 2.6 Data analysis

Data analysis was performed in R project using the packages “dplyr”, “reshape2”, “lme4”, “lmerTest”, “ggplot2”, “ggpubr”, “ggbeeswarm”, “Hmisc”, “corrplot”. The dynamics of the parameters from the heartbeat discriminations task (threshold, slope, and confidence), state anxiety, attention state and heart rate were assessed using mixed linear models with the interacting variables time and group (haptic or visual) as fixed effects and participant ID as a random effect. Post-hoc comparisons were performed using paired Wilcoxon tests, with Bonferroni multiple comparison correction of p-values. Differences between the user ratings of helpfulness and pleasantness of the haptic and visual training were assessed with Mann-Whitney test. Pairwise correlations were explored using Spearman method.

## 3. Results

### 3.1 Training with haptic feedback led to more accurate heart rate discrimination and higher confidence

Training with haptic feedback (see Figure 1,) resulted in a decrease in absolute interoceptive threshold value (towards more accurate estimation) and an increase in confidence in heart rate discrimination (see Tables 1 and 2, Figure 3, panels A and B). At the same time, no effects of time, group or their interaction were observed for interoceptive slope, as well as threshold and slope in the sound rate discrimination condition. Confidence in sound rate discrimination was higher in the main group (effect of group: estimate 8.0, t=2.1, p=.041) and demonstrated a minor decrease after both types of training (effect of time: estimate -3.3, t=1.6, p=.045).

**Figure 3.**
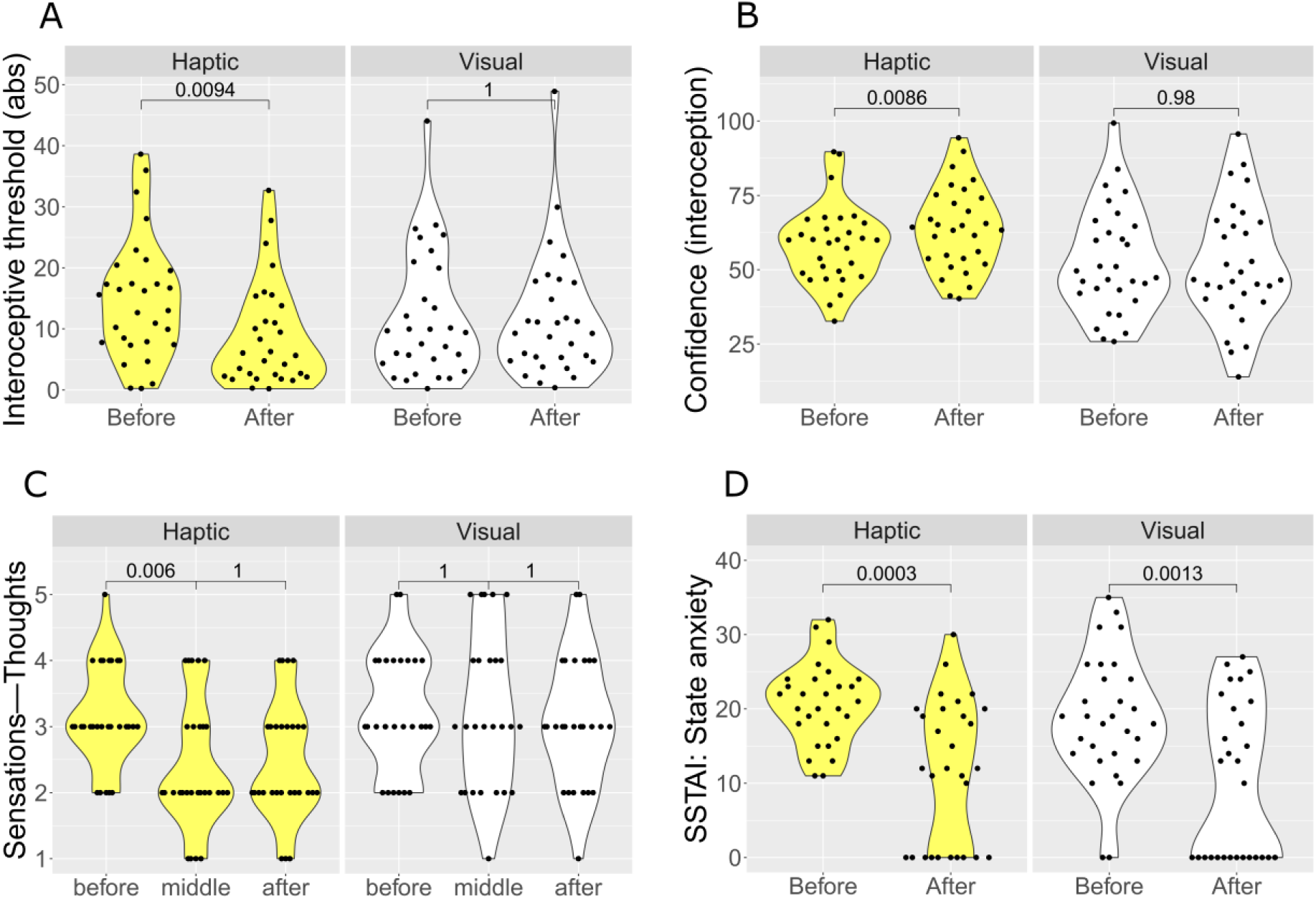
Changes in the interoception and anxiety after the haptic vs. visual heartbeat feedback training. *Note*. Real-time haptic but not visual feedback resulted in decreased interoceptive threshold (indicative of higher interoceptive accuracy), increased confidence in heart rhythm discrimination, and a shift of attention towards sensations. Both trainings led to a decrease in state anxiety level. Indicated are p-values for the post hoc comparisons with paired Wilcoxon tests (Bonferroni-corrected)

**Table 1.**
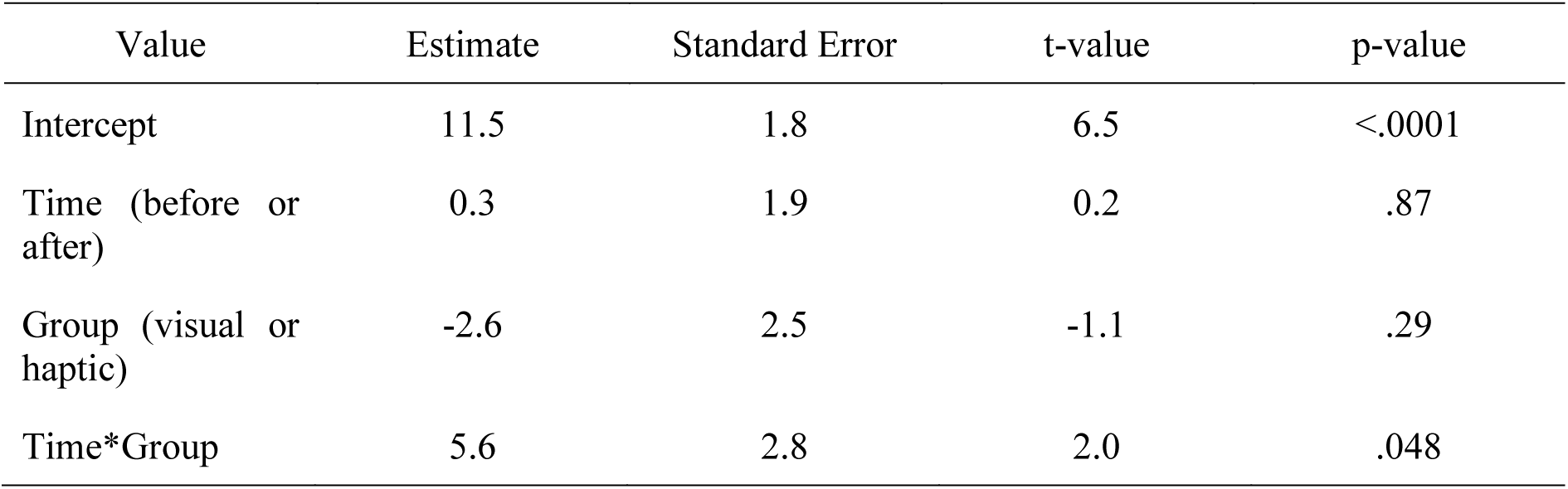
Results of the mixed linear modeling of the influence of training type on the dynamics of interoceptive threshold (absolute value)

| Value | Estimate | Standard Error | t-value | p-value |
| --- | --- | --- | --- | --- |
| Intercept | 11.5 | 1.8 | 6.5 | <.0001 |
| Time (before or after) | 0.3 | 1.9 | 0.2 | .87 |
| Group (visual or haptic) | -2.6 | 2.5 | -1.1 | .29 |
| Time*Group | 5.6 | 2.8 | 2.0 | .048 |

**Table 2.** Results of the mixed linear modeling of the influence of training type on the dynamics of confidence during heart rate discrimination.

| Value | Estimate | Standard Error | t-value | p-value |
| --- | --- | --- | --- | --- |
| Intercept | 51.8 | 3.0 | 17.2 | <.0001 |
| Time (before or after) | 1.0 | 1.6 | 0.7 | .51 |
| Group (visual or haptic) | 12.0 | 4.3 | 2.8 | .006 |
| Time*Group | -6.3 | 2.2 | -2.8 | .007 |

### 3.2 Training with haptic feedback led to a shift in attention towards sensations

Attentional focus shifted towards sensations during the training with haptic feedback, but not during the training with visual feedback (see Figure 3C, Table 3).

**Table 3.**
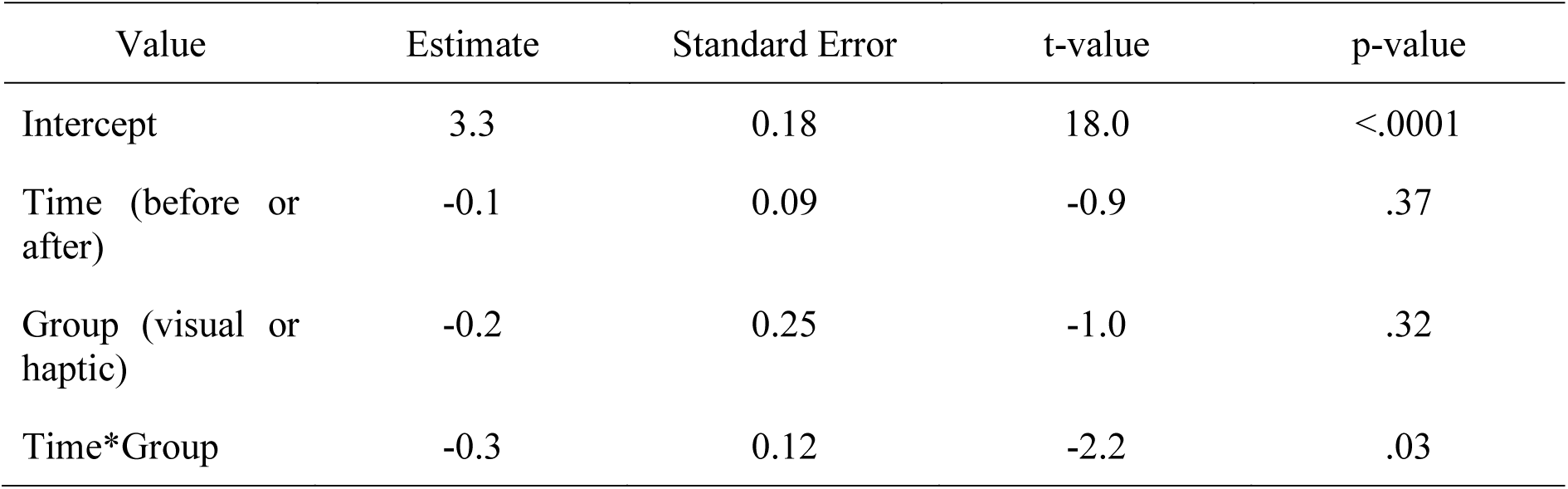
Results of the mixed linear modeling of the influence of training type on dynamics of the attentional focus.

| Value | Estimate | Standard Error | t-value | p-value |
| --- | --- | --- | --- | --- |
| Intercept | 3.3 | 0.18 | 18.0 | <.0001 |
| Time (before or after) | -0.1 | 0.09 | -0.9 | .37 |
| Group (visual or haptic) | -0.2 | 0.25 | -1.0 | .32 |
| Time*Group | -0.3 | 0.12 | -2.2 | .03 |

**Table 4.** Results of the mixed linear modeling of the influence of training type on dynamics of the state anxiety.

| Value | Estimate | Standard Error | t-value | p-value |
| --- | --- | --- | --- | --- |
| Intercept | 9.9 | 1.6 | 6.1 | <.0001 |
| Time (before or after) | 8.8 | 1.9 | 4.6 | <.0001 |
| Group (visual or haptic) | 1.7 | 2.3 | 0.8 | .45 |
| Time*Group | 0.3 | 2.7 | 0.09 | .92 |

### 3.3 Training led to a decrease in anxiety

State anxiety demonstrated a non-specific decline after training of any type (see Figure 3D, Table 3).

### 3.4 Training with haptic feedback was rated as more pleasant

In the debriefing questionnaire, participants rated training with haptic feedback as more pleasant (7.7±1.3 vs. 5.6±2.7, p = .001) and helpful (6.8±1.9 vs. 4.9±2.9, p = .01; see Figure 4).

**Figure 4.**
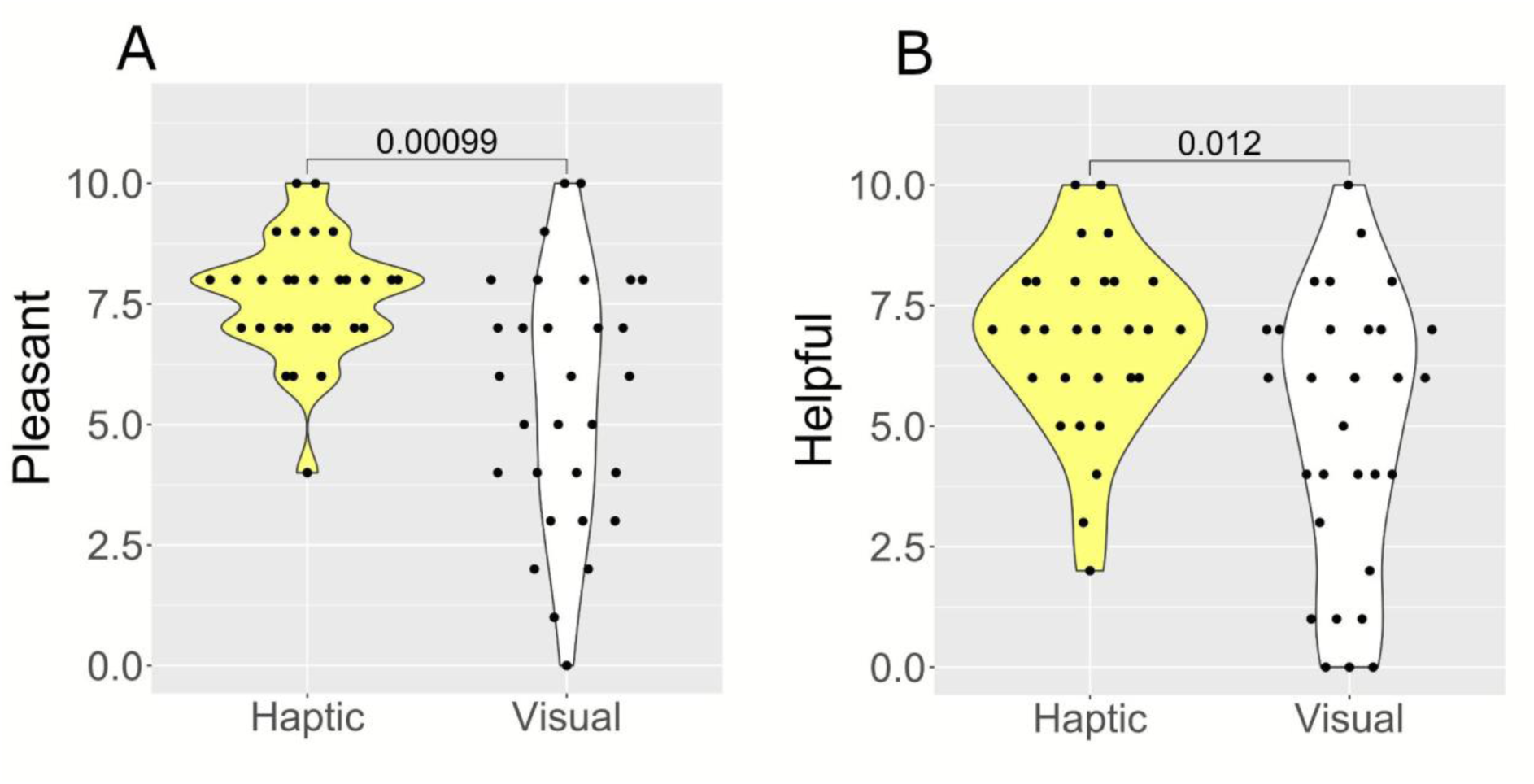
Users’ feedback for the haptic and visual trainings. *Note.* Real-time haptic feedback was rated by users as more helpful and pleasant. Indicated are p-values for the comparisons with Mann-Whitney test.

### 3.5 Interaction between the effects

Pairwise correlations between the changes in interoceptive threshold and confidence, attention state, and state anxiety (after − before), as well as pleasantness and helpfulness ratings were explored to investigate the possible interactions between the effects (see Figure 5). Shift of attention towards the body positively correlated with subjective helpfulness of the training (R = −.29, p = .037; lower attentions ratings correspond to the attention towards the body), and a correlation between helpfulness and pleasantness was observed (R = .41, p = .001).

**Figure 5.**
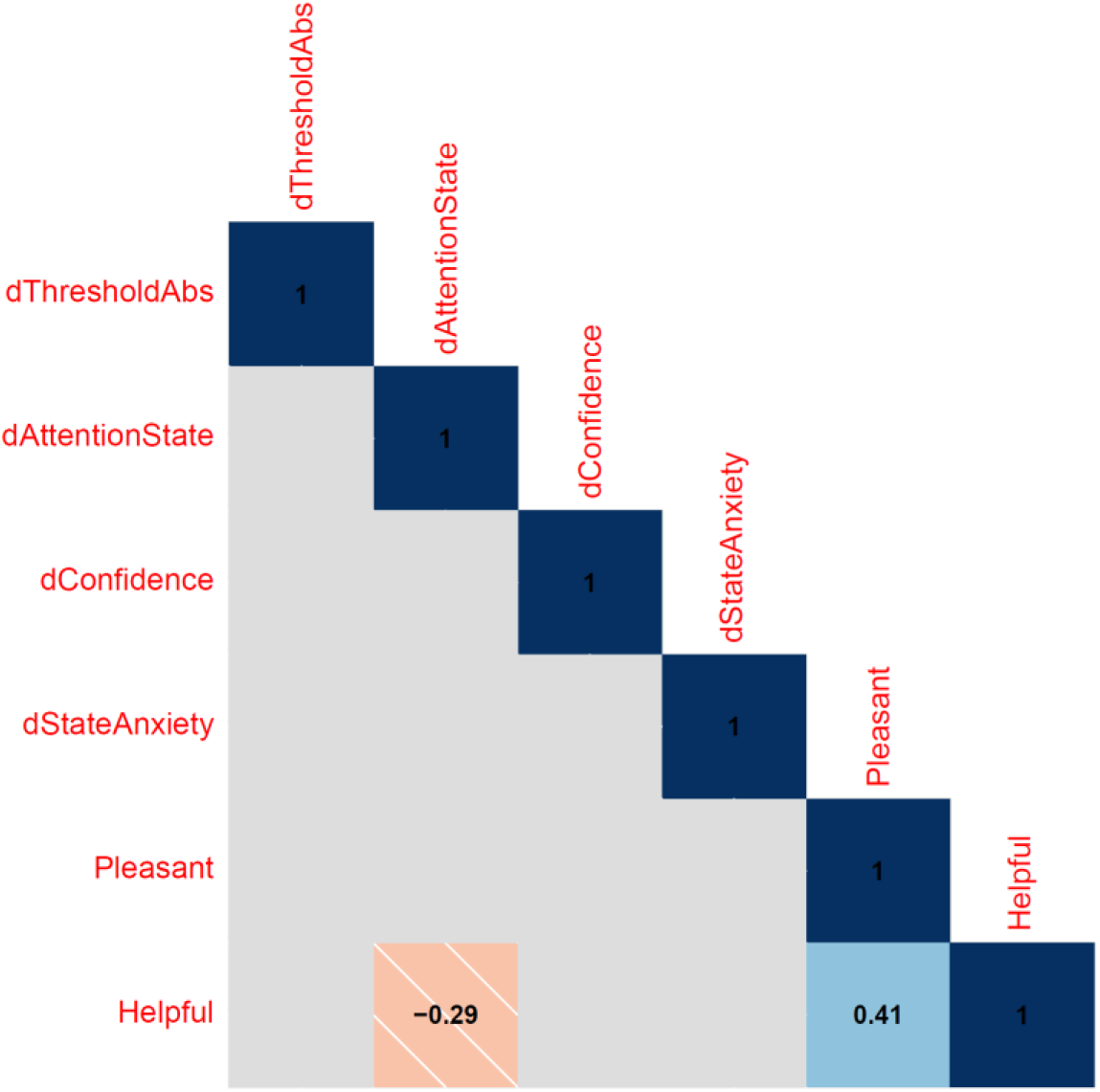
Correlations between the different effects of intervention. *Note*. Subjective ratings of helpfulness and pleasantness were correlated, and helpfulness was additionally associated with a shift of attention towards the body. Other tested correlations did not reach the level of significance. Indicated are correlation coefficients estimated with the use of Spearman test, only significant correlations are presented with p-value threshold of .05.

## 4. Discussion

### 4.1 Summary of key findings – visual vs. haptic real-time feedback of heartbeat

The study demonstrates a positive shift in cardioception after a single session of real-time haptic feedback and shows its superiority over more traditional method of delivering feedback in the visual modality. Following the haptic intervention, increases were observed in different dimensions of interoception (Garfinkel et al., 2016): interoceptive accuracy (as indicated by the threshold of the psychometric function, interoceptive sensibility (represented by confidence), and attention towards the body (see Fig. 3, Tables 1–3). The dynamics in the different dimensions occurred independently, suggesting that haptic feedback exerts parallel effects on multiple levels at the same time (see Fig. 5). The changes were observed only after haptic, but not visual feedback, rendering unlikely the role of unspecific factors such as learning the heart rate, habituation to the heart rate discriminations task, or relaxation. Furthermore, the lack of change in the sound rhythm discrimination task indicates that our training specifically targets interoception rather than leads to general, modality non-specific improvements in rhythm discrimination or attention. These results support our hypothesis regarding the higher efficacy of naturalistic haptic vs. visual feedback in targeting interoception. In addition, the developed intervention might be superior in terms of user experience: it was perceived as more pleasant and helpful than training with visual feedback.

### 4.2 Implications for sensory research

The observed beneficial effects of haptic feedback modality align with results of our previous studies demonstrating utility of haptic supplementation of audial information. For example, it has been shown that haptic feedback of speech leads to an increase in speech comprehension in noisy environments of 6db with no training at all (Cieśla et al., 2019). With very limited training, this rises to an increase of 10db (Cieśla et al., 2022), providing a great improvement to the ability to comprehend speech in difficult settings. In addition, speech supplemented with haptic feedback was found to be more persuasive (Saint-Aubert et al., 2023). Thus, tactile channel can be effectively used to support other sensory modalities, and the current study expands the previous evidence to the field of interoception, where such approach is even more promising.

In contrast with previous studies (Meyerholz et al., 2019; Schillings et al., 2022), we did not observe a change in interoceptive abilities after a training with visual feedback only. This finding probably originates from a much shorter duration of the training—12 min in our study vs. longer training in studies focused on visual feedback. The participants might not have enough time to build the connection between less intuitive visual feedback and inner sensations from the heart. Haptic real-time feedback appeared to be effective even in such a brief setting making it more suitable for both clinical rehab and for general mindfulness purposes.

The observed superiority of naturalistic haptic vs. visual feedback can also be explained within the context of the concept and theory of predictive coding (Apps & Tsakiris, 2014; Owens et al., 2018). During perception, incoming sensory information is aligned with a priori models. When it comes to cardioception, knowing how the heartbeats feel allows recognition of the relevant interoceptive information within the bottom-up stream. People with low interoceptive abilities might struggle to filter in the sensations originating from the heart due to a lack of an a priori model. When provided with feedback, they receive an external template with which to match inner sensations. In the case of visual feedback, this external template only includes the timings of the beats. Haptic feedback contains additional information about where and how the beats might be felt, thus representing a much richer cue. External haptic stimuli matched with inner sensations in time, location and waveform might serve as excellent guidance for finding the interoceptive sensations originating from the beating heart.

The importance of aligning artificial stimuli characteristics with the natural bodily prototype has been repeatedly demonstrated on the rubber hand illusion model (Ehrsson, 2019). During the rubber hand illusion experiment, an artificial hand is positioned in front of the participant, while their real hand is hidden. Synchronous stimulation of the real and rubber hands with a brush results in a sense of ownership over the rubber hand (embodiment). The strength of the illusion depends on the similarity between the characteristics of the real and rubber hands, such as position (Ide, 2013), size (Pavani & Zampini, 2007) and color (Lira et al., 2017). Thus, in terms of predictive coding, congruent stimuli are more likely to be incorporated into perceptual models of the body (Dobrushina et al., 2021; Reader & Crucianelli, 2019). We propose that in our setup the similarity between delivered haptic stimuli and natural heartbeat sensations might support feedback embodiment with enhancement of inner models underlying interoception. This hypothesis aligns with the previously described efficacy of cardio-visual full body illusion in which providing exteroceptive information related to the heartbeats by illuminating a silhouette induces self-identification with the virtual body (Heydrich et al., 2018).

### Implications for neurotech, limitations and future directions

The method, technology, and approach we have studied are promising as a new opportunity for health tech, neurowellness, and technologically enhanced mindfulness applications. Our results indicate that haptic feedback helps to focus on the heartbeats in an implicit and non-demanding way, simply attracting the attention to the intensified sensations. Thus, it might require less cognitive effort than conventional feedback training and may also be more ecologically sound, since naturally interoception is accomplished without the use of visual inputs but rather with directing the attention to one’s own body. Coupled with high user satisfaction, this makes the developed method attractive for utilization in patients with mental and somatic disorders linked to altered interoception and for people willing to improve bodily and emotional regulation. This list of potential applications includes treatment of depression, anxiety disorders, posttraumatic stress disorder, somatic symptom disorder, borderline personality disorder, improving the quality of neurodiverse people’s lives, enhancing emotional intelligence, and soft skills training (Brewer et al., 2021; Nord & Garfinkel, 2022). Another potential use of our setup is controlled interoceptive exposure in panic disorder (Lee et al., 2006): regulating the intensity of vibration, it is possible to induce mild to strong heart-like sensation.

A modified setup combining two or more haptic feedback systems can be employed for modulating interpersonal interaction, enhancement thereof and increased social relationships, or alternatively for decreasing social pain and isolation (J. Werner et al., 2008). This approach relies on the principle of biobehavioral synchrony (Feldman, 2012): attachment and related oxytocin release involve synchronization in physiological rhythms, including heartbeats (Atzil et al., 2012; Feldman, 2012). Our method could be utilized for providing a haptic representation of another’s heartbeats, which could be particularly effective due to the benefits of tactile stimulation and social touch on cognitive and emotional wellbeing. Another feasible modification is integration of our approach with other technologies. Many systems such as mobile phones, smart watches, virtual, augmented, and extended reality and gaming systems include advanced haptics as well as physiology sensing. Incorporating the developed training or similar one into these available platforms opens interesting possibilities for designing new digital healthcare interventions.

Our study is not free from limitations that highlight the need for further research. We performed a single-session intervention, and only immediate before-after effects were evaluated. Thus, it will be interesting to perform a follow up study with a multi-session study to check the stability of the effects. Since the sample included healthy adults, evaluating the transferability of results to clinical samples remains another target for future studies. A particularly interesting avenue of research is incorporating the setup into behavioral or neuroimaging experiments, allowing exploration of the links between interoception, emotional processing and cognition on the neural level.

### Concluding remarks

The study supports feasibility and superiority of interoceptive training with haptic feedback and extends the existing research (Meyerholz et al., 2019; Quadt et al., 2021; Schaefer et al., 2014; Schillings et al., 2022) by developing a novel method incorporating the principles of biofeedback and sensory augmentation in an accessible setup. We suggest that our approach can be used for enhancement of physical, mental, and social well-being, both as a stand-alone or in integration with other digital health interventions. The results of our study inform the development of mind-body technologies in general, indicating the importance of matching feedback sensory characteristics to the natural bodily signals.

## Author contributions

Olga Dobrushina: Conceptualization, Methodology, Software, Investigation, Formal analysis, Writing - Original Draft, Writing - Review & Editing, Visualization, Project administration. Yossi Tamim: Methodology, Investigation. Iddo Yehoshua Wald: Conceptualization, Methodology, Writing - Original Draft. Amber Maimon: Conceptualization, Writing - Original Draft, Writing - Review & Editing. Amir Amedi: Conceptualization, Writing - Review & Editing, Supervision, Funding acquisition.

## Data and materials availability

The data analyzed during the current study are available from the corresponding author on reasonable request.

